# Aerosol deposition affects water uptake and water loss of beech leaves

**DOI:** 10.1101/2024.12.19.629383

**Authors:** Irmgard Koch, Ansgar Kahmen, Jürgen Burkhardt

## Abstract

The deposition of aerosols on leaves could significantly influence plant-atmosphere-interaction through the formation of very thin aqueous films that allow the transport of liquid water through the stomata. Such films can be formed by deliquescence and dynamic expansion of hygroscopic aerosols (‘hydraulic activation of stomata’). Two processes that may be associated with stomatal liquid water transport are foliar water uptake (FWU) and the contribution of ‘leaky stomata’ to minimum epidermal conductance (g_min_).

We investigated whether ambient aerosols affect FWU and g_min_ of *Fagus sylvatica* seedlings. Plants were grown in ventilated greenhouses with ambient air or filtered, almost aerosol-free air. The g_min_ was determined using leaf drying curves. FWU was investigated gravimetrically and with deuterium-enriched water, starting from different leaf water potentials, by spraying freshly-cut or pre-dried leaves (60 minutes).

The presence of aerosols in the environment increased g_min_ by about 47%, confirming previous measurements in other species. Aerosols also increased FWU measured by deuterium uptake. FWU was higher for freshly-cut leaves than for pre-dried leaves, despite the lower leaf water potential. No gravimetric weight gain could be detected.

Both the g_min_ and FWU results are consistent with bidirectional stomatal transport of liquid water along aerosol-induced pathways. The FWU result could also have been generated by water vapor through ‘reverse transpiration’, although the functional contribution of the aerosols would remain unclear. At low leaf water potential, the pathway may dry out and become less functional for FWU, whereas it may still be noticeable as stomatal leakage, given the strong gradient of water potential from the leaf interior to the atmosphere.

## Introduction

Aerosol particles are an essential and ubiquitous component of the atmosphere. They originate from natural sources such as sea salts, desert rock dust, biological particles like microorganisms and plant matter as well as natural combustion residues from forest fires and volcanic activity (Prather et al. 2008; Hamilton 2015). However, their concentration has increased massively since industrialization (Andreae 2007) primarily due to emissions from fossil fuel combustion and agricultural practices (Heintzenberg 1989; Prather et al. 2008; Hamilton 2015; Burkhardt and Grantz 2017). Between 25 to 50% of the aerosols have hygroscopic properties, enabling them to absorb moisture from the environment (Duce and Tindale 1991; Mahowald et al. 1999; Burkhardt 2010; Bressi et al. 2021) and may also deliquesce and effloresce (recrystallize) due to humidity changes. Deposited particles on leaf surfaces are still retaining their hygroscopic properties. Within the leaf boundary layer and especially close to stomata, high relative humidity occurs on plant surfaces, enabling the formation of highly concentrated salt solutions by deliquescence even on sunny days (Burkhardt and Eiden 1994; Burkhardt and Hunsche 2013). Apart from ionic salts in aerosols, some organic substances on leaf surfaces are hygroscopic and interact with cuticular transport (Tredenick and Farquhar 2021; Tredenick et al. 2022).

Although the ability of water to enter leaves through stomata has long been doubted due to water surface tension (Schönherr and Bukovac 1972), the formation and extension of thin films has ultimately been confirmed by direct experimental evidence from confocal microscopy (Eichert et al. 2008), by videos from environmental scanning electron microscopy (Burkhardt and Pariyar 2014), and by theoretical explanation about the reduced surface tension of highly concentrated solutions of certain ‘chaotropic’ salts/ions from the Hofmeister series (Pegram and Record 2007; Dos Santos and Levin 2010; Burkhardt et al. 2012). In addition, salt solutions experience dynamic salt expansion from humidity fluctuations that cause changing liquid and solid stage of the deliquescent salts, lead to extension and displacement of salt crusts and the generation of branched/dendritic patterns (Qazi et al. 2019; Licsandru et al. 2023). Such extended salt solution films along stomatal walls may then connect with the end of the hydraulic system inside the substomatal cavity, thus shifting this end to the leaf surface, a process called hydraulic activation of stomata (HAS, Burkhardt 2010). Parallel to water vapour transpiration, HAS establishes a second stomatal pathway of water loss, where liquid water exits the stomata as thin films (typically < 0.1 µm), independent of stomatal aperture. In both aerosol amendment and exclusion studies, the presence of aerosols altered the aperture – flux relationship of *Sambucus nigra* (Burkhardt et al. 2001) and faba bean (Grantz et al. 2018) stomata, and increased transpiration and conductance for beech and hydroponic sunflowers (Burkhardt and Pariyar 2016; Burkhardt et al. 2023). Once established, salt solution films that are still in contact with a water source could still be active and act as a wick even when the stomata are almost or even completely closed. This means they might affect those more passive stomatal water fluxes that do not involve CO_2_ uptake such as nocturnal transpiration (Vega et al. 2023), foliar water uptake (FWU), and minimum transpiration (g_min_).

FWU is a historically neglected part of the plants water balance, partly also caused by the surface tension argument. In the last two decades it has been increasingly researched and is meanwhile considered important, as it has been shown to be present and contribute importantly to the water balance of more than 200 species worldwide (Berry et al. 2019). In the FWU process, plants take up water through the leaves under conditions of a water potential gradient that has lower values inside than outside of the leaf, e.g., drought affected leaves experiencing leaf wetting from rain, fog, or dew (Goldsmith et al. 2017; Vesala et al. 2017; Schreel and Steppe 2020a). Coopman et al. (2021) showed in their experiments in mangroves that deliquescent salts on the leaf surface harvested water from the atmosphere leading to an increase in reversed sap flow, indicating top-down rehydration.

Different pathways for foliar water uptake have been suggested: diffusion trough the cuticle (Ketel et al. 1972), the absorption by hydathodes (Martin and von Willert 2000) and trichomes (Ohrui et al. 2007; Schreel et al. 2020), and last but not least stomatal transport, which may either happen as water vapor (‘reverse transpiration’; Vesala et al. 2017; Griffani et al. 2024; Vesala 2024), or as liquid water as a consequence of HAS (Burkhardt 2010; Schreel and Steppe 2020a). Although this pathway has been considered by several researchers, it has so far not explicitly been tested.

Minimum transpiration (g_min_) is a measure for passive water loss to the unsaturated atmosphere and may become a limiting factor for survival because plants cannot avoid it (Muchow and Sinclair 1989). g_min_ depends on stomatal and cuticular traits (Wang et al. 2024) and varies with the environment including altitude and temperature (Duursma et al. 2019). Caused by measured values too high for explanation by cuticular conductance, stomata have generally considered to be major contributors (Kerstiens 1996). At first look this may seem surprising, as the most common method for determination is repeated weighing of detached leaves, where stomata should be closed. Such contributing stomata have thus been called ‘malfunctioning’, ‘leaky’, ‘incontinent’, ‘unhealthy’, or just ‘incomplete stomatal closure’ (Sase et al. 1998; Takamatsu, Sase, and Takada 2001; Jordan and Brodribb 2007; Duursma et al. 2019). While aerosols and HAS have already been recognized as a possible explanation for increased g_min_ (Burkhardt et al. 2018; Grantz et al. 2018), their importance for FWU has not been tested.

In this study, we used an aerosol exclusion experiment to compare FWU and g_min_ of beech leaves grown in ambient or filtered, almost aerosol free air. We quantified FWU by spraying leaves with deuterium-enriched water, for two different starting points of leaf water potential by pre-drying one group. We also investigated FWU gravimetrically. g_min_ was determined from the weight loss of detached leaves.

The following hypotheses were tested:

- *Aerosols are an environmental factor affecting both g_min_ and FWU*

- *Deposited aerosols increase the deuterium uptake (FWU) of beech leaves*

- *FWU is higher for drier leaves i.e. pre-drying leaves reduces water potential and therefore increases FWU*

## Material and Methods

### a) Plant material

Two-year old *Fagus sylvatica* trees were planted in soil (3 L pots) in spring of 2020 before sprouting and grown in a greenhouse with filtered air (FA) in Bonn (50°43’25.7“N 7°05’04.6”E). Air was filtered in this greenhouse with HEPA filters ensuring removal of 99% of ambient aerosols (Grantz et al. 2018). Nine weeks before experiments were conducted, half of the plants were transferred to a greenhouse with ambient air (AA). Air exchange was the same to ensure similar conditions in the greenhouses. The experiments were conducted in 2020 with the exception of a repeat of the g_min_- measurement. This took place in September 2024 on 9 new beech seedlings that had been grown in soil (3 L pots) in the same greenhouses (5 plants in AA, 4 plants in FA) since budbreak.

In the following experiments, leaf area was repeatedly determined. For this purpose, a photo of the leaf laying completely flat on the work surface was taken next to a size standard. Photos were analysed in ImageJ (Image Processing and Analysis in Java, Version 1.53, Wayne Rasband, National Institutes of Health, USA) by determining the number of pixels and converting them into cm^2^ using the size standard.

### b) Determination of particles on leaf surfaces

Plants were grown in greenhouses with (ambient air, AA) and without particles (filtered air, FA) with the assumption that particles would deposit on leaf surface influencing water fluxes. To quantify different amounts of salts on the leaf surface of AA and FA leaves, for ten plants per greenhouse eight leaves per plant were cut off and leaf area was determined. Leaves were washed in 40 ml ultrapure water (MilliQ Gradient A10, Millipore, USA) in an ultrasonic bath (5 min, 30 °C, Bandelin Sonorex super RK 510 H, Bandelelin electronic, Germany) keeping the petioles out of water. Water samples were filtered (0.45 µm) and water was frozen until analysis. pH and conductivity of water were measured (WTW Series inoLab pH/Cond 720, Xylem Inc., USA). Concentration of Na^+^ and K^+^ was determined with flame photometry (ELEX 6261, Eppendorf AG, Germany). Concentration of Mg^2+^ Ions was measured with atomic absorption spectroscopy (AAS 1100B, PerkinElmer Inc., USA) and NH ^+^ was identified with the continuous flow analyzer (Auto Analyzer 3, Bran und Luebbe GmbH, Germany). With an ion chromatography system (Eco IC, Metrohm Inula GmbH, Austria) concentration of Cl^-^, NO ^-^ and SO ^2-^ was measured.

### c) Minimal epidermal conductance (g_min_)

Minimal epidermal conductance was measured on 19 beech seedlings, with nine in FA-greenhouse and ten plants in AA-greenhouse. The experiment was conducted on 10 plants (five plants each greenhouse) in 2020 and repeated on nine plants in 2024 (5 plants in AA and 4 plants in FA). From each plant, 3 leaves were cut off and the petiole was sealed with Paraffin. Leaf surface area was determined and leaves were weighed in one-hour intervals for 8 h (Scale: Max = 120 g, Min, 0.001 g, d = 0.00001 g, e = 0.001 g, explorer EX125M, Ohaus, USA). In between measurements leaves dried under the laboratory fume hood. Temperature and humidity were measured every 10 seconds (Tinytag-logger, TGP-4500, Gemini Data Loggers Ltd, UK). g_min_ was calculated according to (Sack et al. 2003). Mean values for each plant were taken for statistical analysis with ‘Welch’s unequal variances t-test’.

### d) Deuterium measurements

Deuterium measurements were carried out with five beech seedlings per greenhouse from the different treatment groups. These differed in the spray treatment (deuterium oxide water or tap water), and the drying time before this treatment (freshly-cut: 0 minutes; pre-dried: 60 minutes). This resulted in a total of 40 samples (2 greenhouses x 2 spray treatments x 2 drying times x 5 repetitions). For each sample five leaves were cut off to increase amount of extractable water. Then the petioles were sealed with paraffin to prevent water loss. Leaf weight was obtained with a microbalance (Max = 120 g, Min, 0.001 g, d = 0.00001 g, e = 0.001 g, explorer EX125M, Ohaus, USA) and leaf area was determined.

Pre-died leaves were hung vertically for 60 min and illuminated with a light source (PAR: 200 µmol m^-2^ s^-1^; LED, light intensity 10 %, SL 3500-W-J, Photon Systems Instruments spol. s r. o., Czech Republic), to keep the stomata open and induce water loss, then the leaves were weighed again.

Leaves were exposed to spraying with deuterium oxide water (ratio deuterium oxide water to tap water 1:10 000; 99.9 atom% deuterium oxide, Sigma-Aldrich Chemie GmbH, Germany) or tap water as a control, while hanging vertically illuminated by a light source (PAR: 200 µmol m^-2^ s^-1^; LED, light intensity 10 %, SL 3500-W-J, Photon Systems Instruments spol. s r. o., Czech Republic). For 1 h, the leaves were kept wet, i.e. they were re-sprayed with the respective treatment before they could dry. Then they were dried with paper towels and weighed again.

Samples were then stored frozen in 10 ml exetainers (Labco Limited, UK). They were transferred from the lab in Bonn to the Stable Isotope Ecology Lab at the University of Basel. The water of the leaf samples was extracted with a cryogenic distillation apparatus, explained in detail in Newberry et al. (2017). Shortly, samples were heated in extraction vials (2 h 15 min, 90 °C). The water evaporated and was collected in U-tubes submerged in liquid nitrogen. The sample water was analysed for δ^2^H and δ^18^O [‰ VSMOW] with ‘Thermal conversion/Elemental Analyzer’ (TC/EA) and ‘Delta V Plus Isotope ratio mass spectrometer’ (IRMS) through a ‘ConFLO IV Interface’ (Thermo Fischer Scientific, Germany). Instruments are VSMOW/SLAP-scale normalized and calibrated with internal standard.

Long term analyses show accuracy of 0.7 ‰ for δ^2^H and 0.24 ‰ for δ^18^O.

Hydraulic redistribution is expressed as the percentage of the isotopic composition of water in leaves (leaves sprayed with deuterium oxide - δ^2^H_sample_ and with tap water - δ^2^H_control_) and isotopic composition of water in spray (deuterium oxide water - δ^2^H_spray_; tap water - δ^2^H_tab_; formular 1).

FWU_capacity_ is calculated with the difference of the mass before (Mass_fresh_; g) and after spraying (Mass_sub_; g) divided by the leaf area (A; cm^2^; formular 2; Schreel and Steppe 2020b).

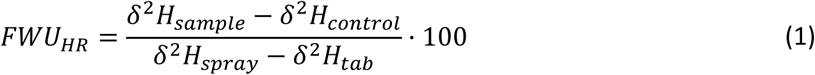

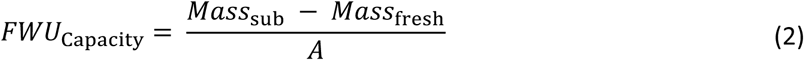

### e) Water potential and leaf conductance

Leaf conductance was measured with a steady state porometer (LI-1600M, LI-COR, USA) on 5 leaves per group. Different leaves but the same five beech seedlings per greenhouse (FA and AA 9 weeks) as in the deuterium experiment were tested. There were two drying times as well (freshly cut: 0 min; and pre-dried: 60 min). Drying took place under the same conditions as the deuterium spraying experiment, with leaves hanging vertically illuminated by a light source (PAR: 200 µmol m^-2^ s^-1^; LED, light intensity 10 %, SL 3500-W-J, Photon Systems Instruments spol. s r. o., Czech Republic). Water potential was measured with a Scholander bomb (Soilmoisture Equipment Coop., USA) on the same leaves as the leaf conductance.

### f) Statistics

g_min_ was tested with ‘Welch’s unequal variances t-test’ because of unequal variances. Washed off ions from the leaf surface were tested with an ‘unpaired two-sample Wilcoxon test’ because data was not normally distributed. Multiple testing was corrected with Holm-Bonferroni method. Water potential, conductance, FWU_HR_ and FWU_capacity_ were analyzed with Analysis of Variance (two-way ANOVA) with post hoc Tukey’s range test for the treatment groups greenhouse (FA and AA) and drying time (0 min, 60 min). All statistical analysis was performed in R (Version 4.4.1).

## Results

The overall mass of the measured ions was 4.53 µg cm^-2^ for beech leaves grown in the ambient air, which was nearly 14 times higher than 0.33 µg cm^-2^ measured on leaves grown in filtered air (Figure 1 a). The strongest reduction of ions to less than 10% was found for sodium and chloride, nitrate and potassium. Sulfate and magnesium were reduced by about one third, similar to overall electrical conductivity (Figure 1 b). The only measured exception was a threefold higher concentration of ammonium on leaves of filtered compared to unfiltered air, which might be related to gaseous transport of ammonia. pH values were not significantly different (Figure 1 c).

**Figure 1:**
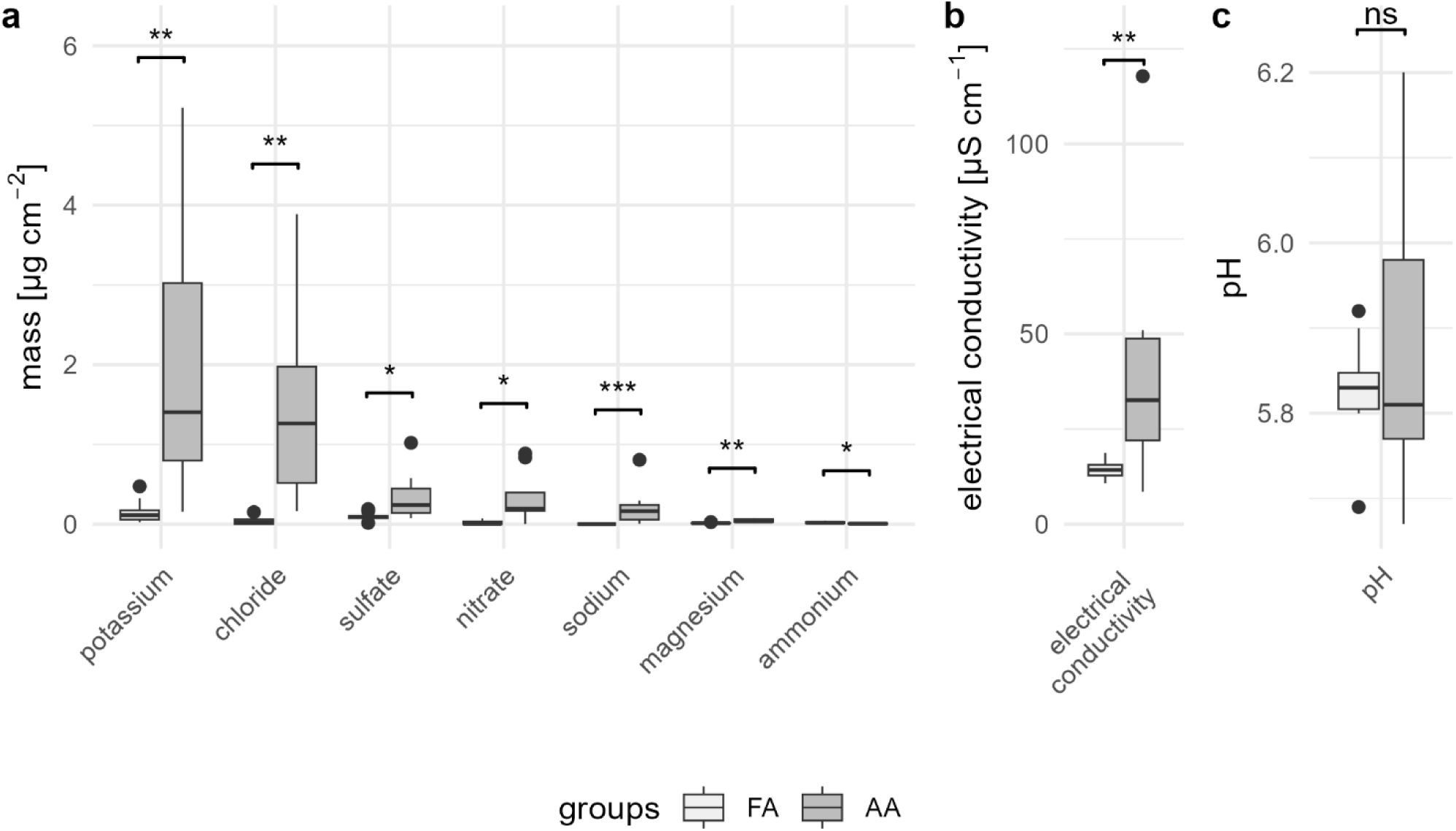
Investigation of washed off water from FA and AA leaves. Masses per leaf area are shown for different ions and elements (a) as well as electrical conductivity (b) and pH (c). Unpaired two-sample Wilcoxon test was used for statistical comparison (0.05 > p ≥ 0.01: *; 0.01 > p ≥ 0.001: **; 0.001 > p: ***; not significant: ns), n=8.

The g_min_ determination in the laboratory showed that leaf surface particles (aerosols) caused faster water loss of detached leaves with time, with significant differences and about 47% higher g_min_ for AA compared to FA leaves (Figure 2).

**Figure 2:**
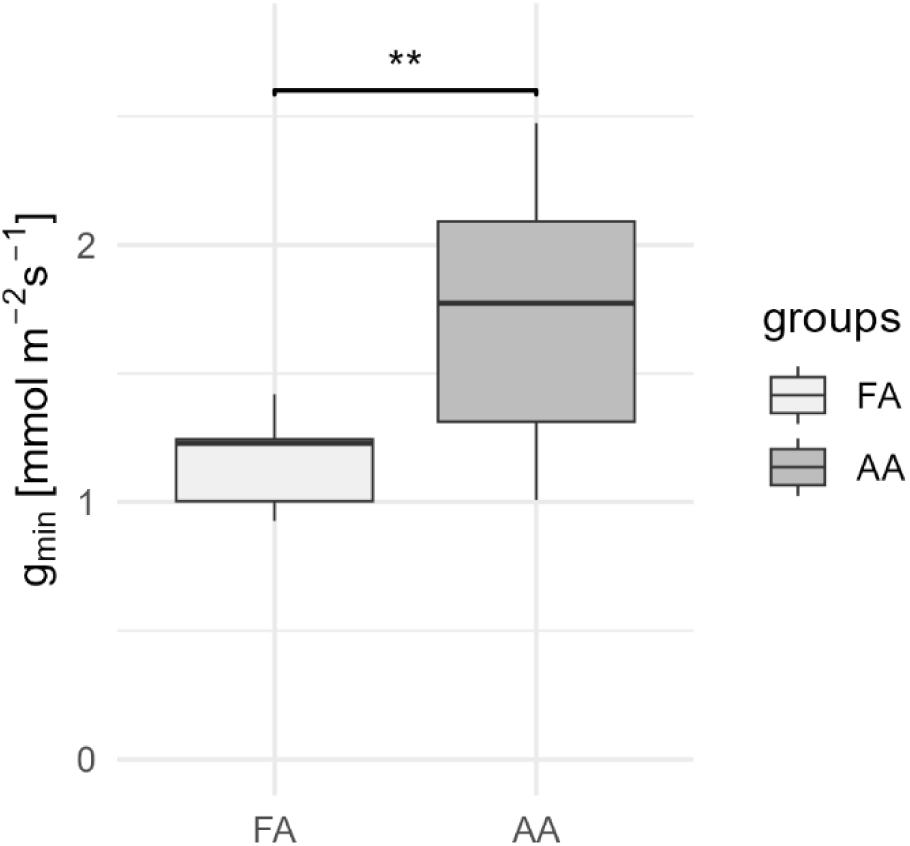
Minimal epidermal conductance (g_min_) for FA and AA mean. Significant difference tested with ‘Welch’s unequal variances t-test’ (p = 0.007 - **), FA: n = 9; AA: n = 10.

The water potential of pre-dried leaves (60 min) was significantly lower by about 1 MPa compared to freshly-cut leaves (0 min) for both greenhouse groups (FA, AA), with considerable scatter. Water potentials were not different between AA and FA treatments for freshly-cut leaves (0 min), but were lower for AA compared to FA leaves for pre-dried leaves (60 min; Figure 3 a).

**Figure 3:**
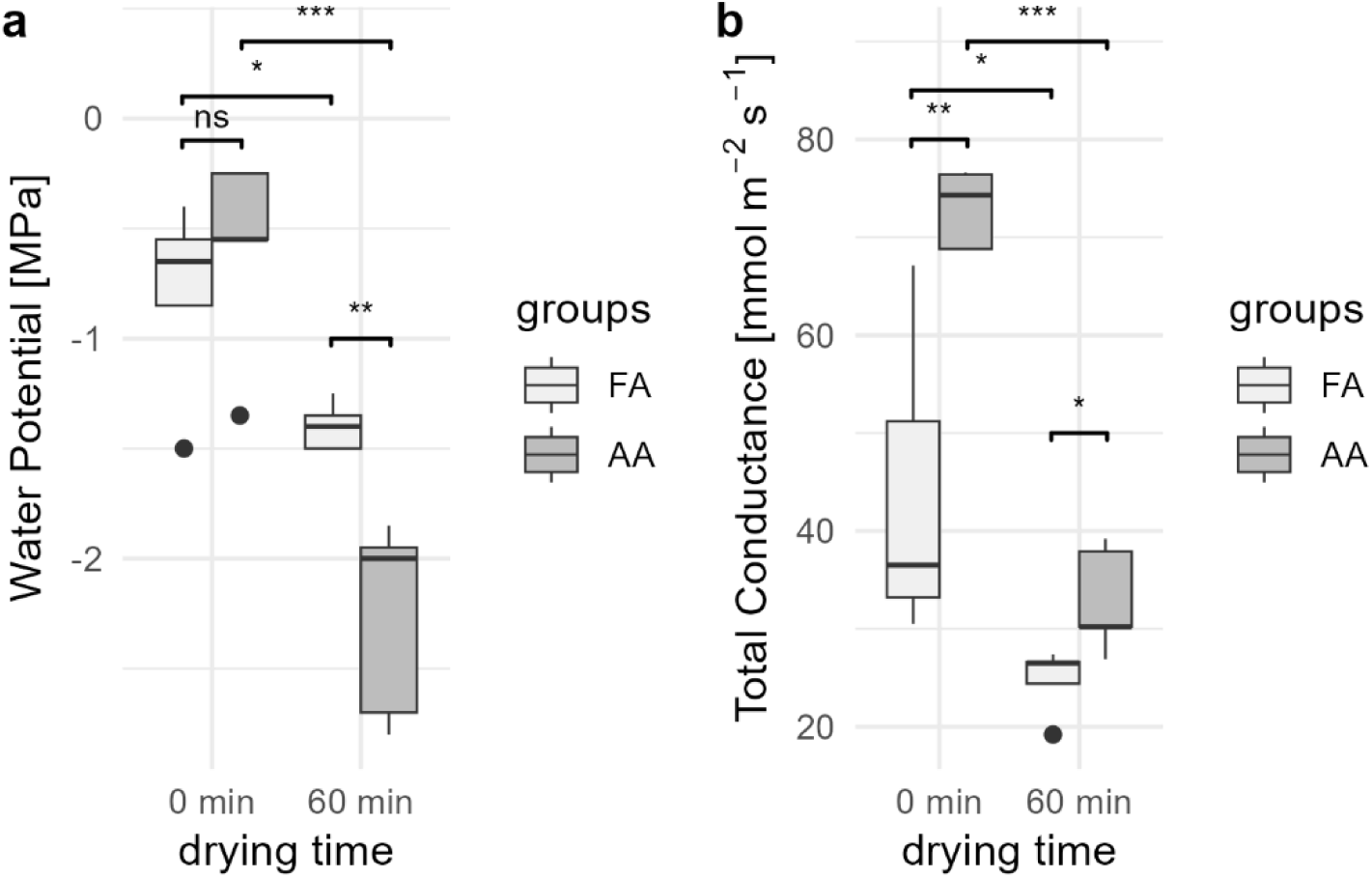
Water potential (a) and total conductance (b) for freshly-cut (0 min) and pre- dried (60 min) samples as well as greenhouse groups FA and AA. Anova with post hoc Tukey test was used for statistical comparison (0.05 > p ≥ 0.01: *; 0.01 > p ≥ 0.001: **; 0.001 > p: ***; not significant: ns), n=5.

The total conductance determined with the porometer were related to the water potentials. They were higher for the freshly-cut leaves (0 min) compared to the pre-dried (60 min) leaves, and in both cases showed higher conductance in ambient air compared to filtered air (Figure 3 b).

FWU_HR_ calculated from deuterium labels showed significantly higher uptake for seedlings grown in ambient air (AA) compared to filtered air (FA; Figure 4 a). There was no significant interaction effect between greenhouse and drying time (F (1,15) = 2.5, p = 0.13), so the main effects were analysed (greenhouse: F (1,15) = 23.5, p = 0.0002; drying time: F (1,15) = 13.4, p = 0.002). For pre-dried leaves (60 min) deuterium uptake was significantly lower than for freshly-cut leaves (0 min; Figure 4 b).

**Figure 4:**
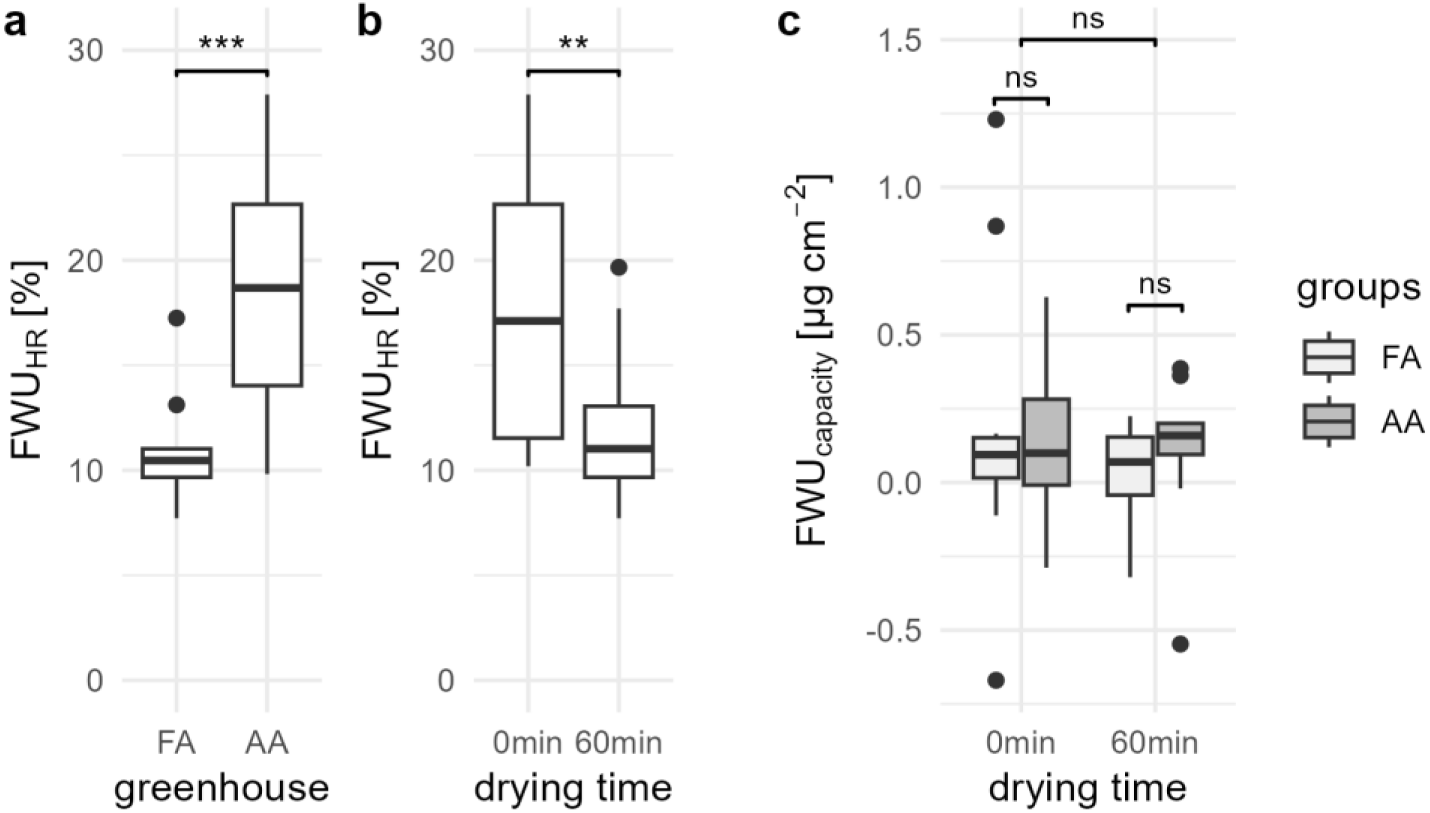
FWU_HR_ is shown for the two groups greenhouse (a) and drying time (b). Two- way anova showed no significant interaction effects so main effects are shown here in two plots. FWU_capacity_ (c) is shown for greenhouse groups (FA, AA) and drying time (0 min – freshly cut, 60 min – pre-dried). Significant differences were tested with two- way Anova (0.05 > p ≥ 0.01: *; 0.01 > p ≥ 0.001: **; 0.001 > p: ***; not significant: ns).

Weighing (FWU_capacity_; Figure 4 c) showed no significant differences in water uptake of groups (greenhouse: F (1,36) = 0.03, p = 0.8; drying time: F (1,36) = 0.7, p = 0.3; Interaction (greenhouse*drying time): F (1,36) = 0.3, p = 0.6). In each group (n = 10) we found two to four samples with a weight loss instead of weight gain.

## Discussion

The surface of plant leaves is not clean. Under normal conditions, they continuously collect and accumulate aerosols. Sub-micron aerosols, which are the most hygroscopic ones, react very little to gravity, but are more influenced by leaf surface micro-roughness, which is one of the reasons why they are not easily washed down. Another reason is deliquescence and the formation of salt crusts, which may create persistent structures on leaf surfaces. Many deposited aerosols thus lose their singular crystalline structure, which makes them part of salt crusts, and can make them indistinguishable from waxes in SEM images (Burkhardt et al. 2018).

In this study, we widely excluded aerosol deposition in the FA-greenhouse by filtering out about 98% of all particles with H13 HEPA filters. The so established 50-fold airborne aerosol concentration difference between the two greenhouses translated into a 14-fold mass difference found on leaf surfaces in the two greenhouses as determined by the washing procedure. Possible reasons for this difference in airborne aerosol concentration and deposition on leaves might mean that even intensive leaf washing did not dissolve all deposited aerosols from the leaves, or that leaching from the inside of the leaves contributed. The amounts of aerosols washed from beech leaves were smaller than the ones previously washed from fir needles (Burkhardt et al. 2018, Table 3). In filtered air (FA), the overall amount on beech was 0.3 µg cm^-2^, while filtered air values in fir had been 1.7 µg cm^-2^ in 1-year-old and 5.0 µg cm^-2^ in 2-year-old needles. In ambient air (AA), 4.5 µg cm^-2^ were found on beech leaves now, similar to earlier values of 5.6 µg cm^-2^ for 1-year old fir needles, and less than 9.0 µg cm^-2^ for 2-year-old fir needles (Burkhardt et al. 2018). Different from the earlier study, the chemical composition was dominated this time by aerosols K, Cl, and sulfate, while almost no ammonium was found. The fact that ammonium was higher on leaf surfaces in filtered than in ambient air might perhaps be due to differences in the phyllosphere of leaves.

The overall g_min_ values were consistent with the ones reported earlier for beech leaves (Kerstiens 1996; Wang et al. 2024). The AA plants showed 47% higher g_min_ than FA plants, showing a faster rate of drying out. This confirms other aerosol exclusion studies with *Quercus robur*, *Pinus sylvestris*, and *Abies alba* (Burkhardt et al. 2018), as well as *Vicia faba* (Grantz et al. 2018), where g_min_ was also lower in filtered air compared to ambient air. Spraying with different 50 mM salt solutions increased g_min_ of Scots pine needles by 18% (sodium nitrate), 34% (sodium chloride and ammonium sulfate), 45% (potassium iodide, a chaotropic salt), and 96% (NaCl with additional surfactant; Burkhardt and Pariyar 2014). In addition to the factors considered by Duursma et al. (2019), aerosol deposition in general can thus be considered an environmental factor increasing minimum transpiration, most likely by increasing the stomatal contribution to g_min_.

It is generally recognized that the stomatal contribution forms a significant part of g_min_. Different from FWU, so far contributions of trichomes and hydathodes do not seem to have been discussed. Stomata probably do not have a foreseen function in this water loss, so stomatal contributions do seem more like an accident, which is also notable from the names that have been given to this phenomenon: ‘malfunctioning’, ‘leaky’, ‘incompletely closed’, ‘incontinent’ (Sase et al. 1998; Takamatsu, Sase, and Takada 2001; Jordan and Brodribb 2007; Duursma et al. 2019).

‘Wax degradation’ is correlated with decreased water potential (Huttunen et al. 1981) and with increased minimum epidermal conductance (g_min_) (Cape and Fowler 1981; van Gardingen et al. 1991; Heinsoo and Koppel 1998; Sase et al. 1998; Anfodillo et al. 2002). Numerous indications also found that g_min_ increases with air pollution particularly connecting ‘wax degradation/erosion’ with the two as well (Cape and Percy 1996; Takamatsu, Sase, and Takada 2001). Takamatsu, Sase, and Takada (2001) found an increasing g_min_ with the ageing of Cryptomeria needles (Takamatsu, Sase, and Takada 2001; Takamatsu, Sase, Takada, et al. 2001) together with increasing ‘wax erosion’ and assumed a ‘Clogging of stomata’ by aerosols (Takamatsu, Sase, and Takada 2001). The appearance of wax erosions might have been misinterpreted in the past and could be a sign of salt crusts resulting from deposited aerosols instead (Burkhardt and Pariyar 2014; Burkhardt et al. 2018).

Roth-Nebelsick et al. (2023) recently measured the weight gain by FWU for seven Pinus species, and found strongest uptake in older needles, which increased when a surfactant was used. They also noticed higher ‘wax erosion’ in the stomatal regions of old needles. Because Pinus species do have neither trichomes nor hydathodes, this is probably the first study where FWU could be tracked to stomatal uptake. For beech, the study situation is very different. Schreel and Steppe (2020a), using several fluorescent tracers and also micro-CT, found clear evidence for almost the entire FWU happening by trichomes, and some very small cuticular uptake, but did not find any stomatal uptake and thus excluded any stomatal contribution for *Fagus sylvatica*. However, in their experiment, stomatal uptake might have been prevented by the experimental limitations, caused by the way in which the tracer solutions were applied: microcentrifuge tubes containing the tracer solutions were attached to the leaves in a way that the tracer solution was in full contact with the leaf surface, and tubes were covered with aluminium foil in order to avoid photodegradation (Schreel and Steppe 2020a). The continuous full contact of the solution with the leaf surface brings up the water surface tension argument, according to which pressure would be needed to force solution into the stomata geometric architecture with its widening substomatal cavity (Schönherr and Bukovac 1972).

Additionally, restricting the access of light by the aluminium foil made stomata probably close. FWU by HAS and the stomatal pathway can thus not be fully excluded for beech leaves by the experiment from Schreel and Steppe (2020a).

A significant increase of FWU_HR_ calculated from the deuterium label uptake was observed in leaves grown in ambient compared to filtered air (Figure 4). Values were significantly higher in freshly-cut leaves compared to pre-dried ones, but we could not show an increase of weight gravimetrically between greenhouse treatments or freshly-cut and pre-dried leaves. We agree with Goldsmith’s statement (Goldsmith et al. 2017) that changes in isotope ratios only show water fluxes and to show FWU as a net increase in mass of water in the leaf, data must be supported by gravimetrical measured net weight gain. The deuterium uptake in our experiment shows that aerosols reduced the resistance for water transport into the leaf. Separately from the aerosol amount, the deuterium uptake was lower for pre-dried leaves although water potential was also lower, which could be indicating better conditions for FWU. A probable explanation is that the salt solution bridges causing HAS were falling dry by the little availability of water for the pre-dried group. Such dried salt crusts have less capacity for water transfer compared to the wet state (Licsandru et al. 2023). A lower influence of HAS on transpiration with increasing VPD was also observed in hydroponic sunflowers (Burkhardt et al. 2023).

The fact that we found FWU of deuterium labelled water triggered by the presence of aerosols does not exclude any of the proposed pathways, so it might have happened via trichomes, the cuticle, and gaseous or liquid passage through the stomata (hydathodes do not seem to be present on beech leaves; Schreel and Steppe 2020a). However, a functional importance of deposited aerosols is most likely for liquid stomatal uptake along aerosol-induced thin aqueous films due to HAS and into trichomes, where they may fulfil a similar function for water entering into the cellulose-rich non- lignified cell walls of midvein trichomes on beech leaves (Schreel and Steppe 2020a). Although asymmetric structure and cellulose components of cuticles may provide preferential water uptake pathways (Kamtsikakis et al. 2021; Tredenick and Farquhar 2021), the strong water attraction of salts together with the low cuticular conductance is probably disadvantageous for FWU. And there is also no obvious mechanistic function of deposited aerosols in the postulated ‘reverse transpiration’ process.

‘Reverse transpiration’, i.e. gaseous uptake of deuterium enriched water, cannot be excluded as a possible pathway leading to the reported results. The conductance of water vapour transport consists of two parts: the stomatal and the boundary layer conductance. The dominant stomatal part is determined by stomatal aperture. It is usually measured with a porometer as was the case here.

Such devices measure the total conductance of the leaf within the cuvette, but usually the results are interpreted as stomatal conductance, because ventilation within the cuvette minimizes the boundary layer resistance, and the cuticular contribution is considered negligible. However, stomatal liquid water transpiration resulting from HAS can contribute significantly (Grantz et al. 2018; Burkhardt et al. 2023), and the aerosol impact cannot be isolated by conventional gas exchange or porometric measurements alone because vapor and liquid flux streams through the stomata merge at the site of measurement (Grantz et al. 2018). Therefore, the higher total conductance of ambient air compared to filtered air, which correlated with higher deuterium uptake cannot be interpreted as stomatal conductance, so does not indicate ‘reverse transpiration’ of water vapour. For the same reason, lower total conductance of pre-dried compared to freshly-cut leaves does neither support nor exclude ‘reverse transpiration’, as it does not necessarily mean that stomatal aperture was lower, even if leaf water potentials would suggest this. Possibly, aerosols entering the stomata would decrease the water potential in the apoplast, which could support ‘reverse transpiration’. Capillary condensation and accumulation of salts in the apoplast reduces water potential by reducing saturation vapour pressure, which could be one of the drivers of ‘reverse transpiration’ (Eiden et al. 1994; Vesala et al. 2017; Vesala 2024). However, water potential gradients would drive both water vapour and liquid water transport. A mechanistic explanation for increasing ‘reverse transpiration’ with increasing aerosol concentration would have to explain how aerosols increase water vapour conductance.

A more detailed experiment on the differences between astomatous adaxial and stomatal abaxial flows could help here.

On the other hand, FWU by trichomes cannot be ruled out by our measurements and might even be the most important point of entrance given recent results (Schreel and Steppe 2020a). Different from ‘reverse transpiration’, deposited aerosols might help in bridging gaps of air-filled, non-hydraulic spaces by establishing a thin aqueous water layer.

## Conclusion

In our study we show that aerosols influence water exchange through the leaf surface. On the one hand minimum epidermal conductance increased with aerosol amounts showing an increased uncontrolled water loss through the leaf surface. On the other hand, deuterium label uptake increased with aerosols showing a reduced resistance for water transport into the leaf. Foliar water uptake could not be supported gravimetrically. Different pathways have been discussed in this paper including trichomes, the cuticle and transport through the stomata in gaseous or liquid form but the transport of liquid water along stomata according to the HAS hypothesis might be to date the best explanation for the shown differences in water uptake and water loss in beech seedlings in dependence to aerosols.

## Acknowledgements

The authors would like to acknowledge Angelika Glogau, Lilli Wittmaier, and Daniel Nelson for excellent lab work.

## Authors Contribution

AK and JB developed the research design and methodologies. IK conducted the experiment with inputs from JB. IK performed data analysis with inputs from JB and AK. IK wrote the original draft. JB and AK contributed with review and editing. All authors have read and agreed to the submitted version of the manuscript.

## Supplementary data

### Funding

JB was funded by the Deutsche Forschungsgemeinschaft (DFG, German Research Foundation) – project number 446535617.

### Data Availability Statement

The data underlying this article will be shared on reasonable request to the corresponding author.

